# Reversible RNA Acylating Reagents with Nitro Reduction Strategy

**DOI:** 10.64898/2026.02.04.703650

**Authors:** Yu Hong, Kaizheng Liu, Avleen K. Chawla, Christina-Panagiota Tsingi, Chenkai Yao, Anna M Kietrys

## Abstract

We developed a series of nitro reduction-reversible acylating reagents. Following optimization of the acylation conditions, these reagents were tested for deacylation with sodium dithionite *in vitro*. We applied this reversible acylation to modulate RNAzyme-mediated pre-tRNA maturation, demonstrating its ability to regulate RNA-RNA interactions. Furthermore, the *in vitro* reversible acylation of EGFP mRNA indicated effective control of its translational activity. To explore cellular applications, we validated NQO1-mediated deacylation *in vitro* and then induced hypoxia in HepG2 cells using cobalt chloride, thereby reactivating the function of acylated EGFP mRNA via endogenous NQO1. Overall, this study highlights the potential for developing nitro reduction-reversible acylation as a new strategy for RNA functional control and RNA-based drug modification.

---

RNA-based therapeutics are promising next-generation drugs for cancer, infectious diseases, and neurological disorders.^1-3^

However, they face significant barriers in large-scale applications. For instance, RNA drugs suffer from off-target effects that impact treatment efficiency and cause immunological side effects.^1^ The selectivity of RNA drugs can be improved through ligand conjugations, such as GalNAc, which specifically enables targeted delivery to the liver.^4^ Alternatively, off-target effects can be reduced using stimuli-cleavable modifications.^5^ For example, incorporating a beta-galactosidase-responsive modification into nucleobases allows the oligonucleotide strand to become activatable in the presence of the enzyme.^6^ Recent studies have demonstrated that post-synthetic acylation of the RNA 2’OH group serves as a convenient tool for regulating RNA function, inspiring the integration of intracellular stimuli-cleavable structures into reversible acylation strategies.^7-9^ Although acylations responsive to esterases and thiols have been reported, expanding the range of intracellular stimuli could further enhance therapeutic efficacy and expand the applicability.^7, 10, 11^

Hypoxia, a low-oxygen condition commonly observed within tumors, arises from the uncontrolled growth of tumor cells outpacing blood vessel development, distinguishing cancerous tissue from healthy tissue.^12^ As a response to hypoxia, cancer cells frequently upregulate NAD(P)H:quinone oxidoreductase 1 (NQO1), a cytosolic flavoprotein enzyme that serves as a biomarker of carcinogen metabolism and tumor progression.^13-16^ Classical studies first characterized nitro reduction in bacteria, where nitroreductase from *E. coli* catalyzes the reduction of nitro groups on aromatic xenobiotics. In mammalian cells, NQO1 and related flavoenzymes can fulfill an analogous role, functioning as nitroreductase that reduces aromatic nitro groups via two-electron transfer.^17-19^ However, NQO1 has not been extensively exploited as an intracellular stimulus for RNA-based drugs. In this study, we focused on developing a reversible acylation strategy responsive to nitro reduction. For *in vitro* applications, the reduction is effectively achieved using sodium dithionite (Na_2_S_2_O_4_), while for *in cellulo* acylation, we utilized NQO1 that was overexpressed under chemically induced hypoxia.^20, 21^ We hypothesized that a nitro reduction-reversible acylation strategy would enable the behavior of acylated RNA to differ across cell lines, thereby achieving cell-specific therapeutic effects.

We synthesized the three acylating reagents and prepared stock solutions by mixing nitrobenzyl alcohol derivatives and carbonyldiimidazole in dry DMSO under anhydrous conditions (Scheme S1). The carbonyl imidazole moiety showed high reactivity toward RNA 2’-OH in other acylating reagents reported before.^8, 9, 22, 23^ Each acylating reagent contained a nitro group that could be reduced to an amino group by Na_2_S_2_O_4_ or cellular NQO1 under hypoxic conditions (Figure 1A and 1B). Upon the nitro reduction, 1,6-elimination was triggered to restore the RNA 2’-OH. We introduced different substituents on the aromatic ring for these acylating reagents. The **4N2CIm** bears the -Cl as an electron-withdrawing group, while the **4N3MIm** bears -OMe as an electron-donating group. We anticipated that the different substituents would influence both the acylation and deacylation reactivity. To verify the acylating ability of reagents towards 2′-OH of RNA, we performed the acylation screening with a short oligonucleotide consisting of 21 nucleotides (N_21_). After the 20 ng/µL N_21_ RNA was incubated with 240 mM acylation reagents in 100 mM MES buffer (pH 7.5) at different temperatures, the reaction was analyzed by 30% denaturing PAGE (dPAGE). Acylated RNA strands would gain mass, therefore traveling more slowly in the gel and appearing as upper-shifted bands. N_21_ oligos were effectively acylated by all three reagents at the tested temperatures, as reflected by the upshifted bands produced above the unacylated strands. From the gel image (Figure 1C and Figure S1), we noticed that all of the acylating reagents showed higher reactivity with increasing temperatures. We then quantitatively compared their reactivities at 50 °C by calculating the acylation percentage and average number of acylations per strand by band intensities. **4N2CIm** acylated 90.1% RNA with an average of 2.0 acylated hydroxyls per strand, and **4NIm** acylated 85.2% RNA with an average of 1.7 acylated hydroxyls per strand (Figure 1E). **4N3MIm** was the least reactive among the three, possibly due to its lower solubility; it only acylated 75.6% RNA with an average of 1.4 acylations per strand; thus, it was ruled out in the following experiments.

**Figure 1.**
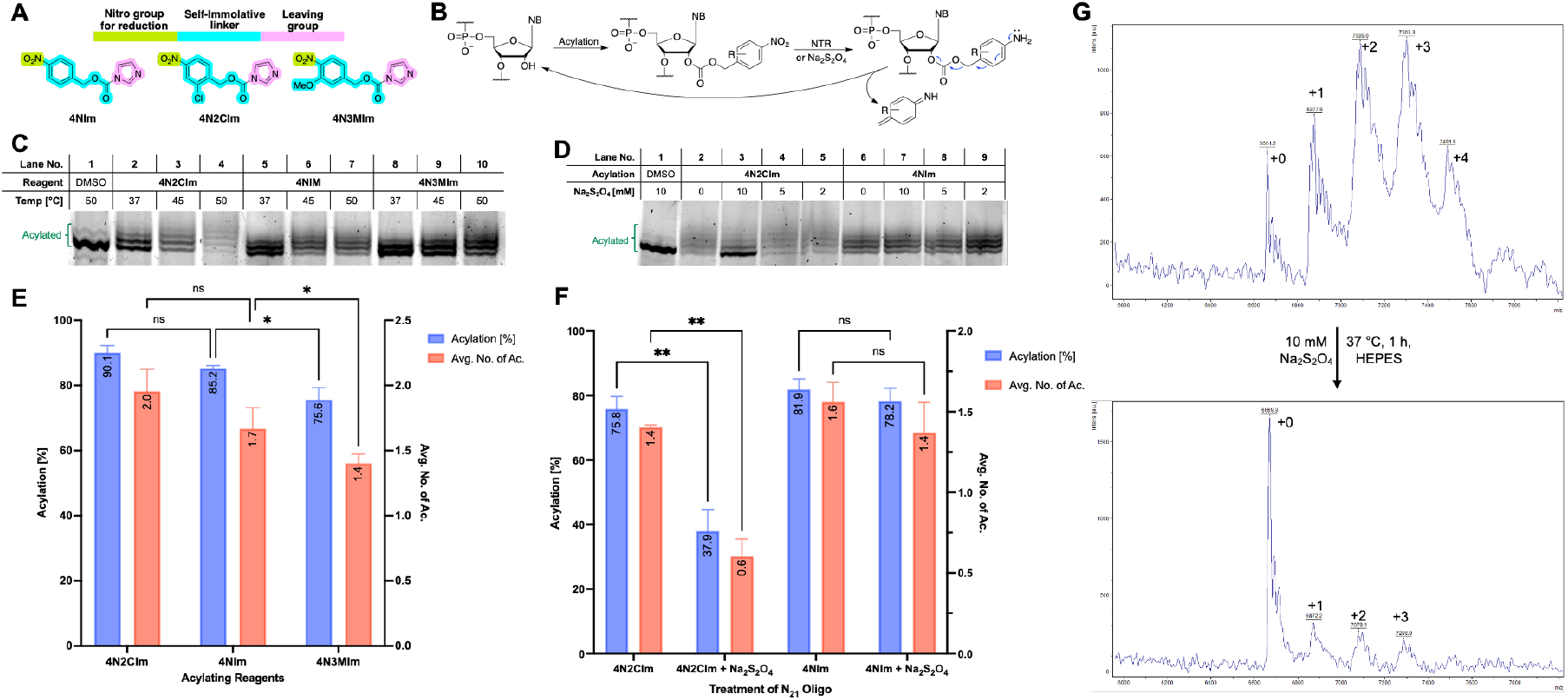
Validation of acylation and deacylation. (A) Design of acylating reagents. (B) Proposed mechanism of reversible acylation via nitro reduction. (C) PAGE analysis: acylation of 20 ng/µL N_21_ oligonucleotide by three acylating reagents (240 mM) at 37, 45, and 50 °C in 100 mM MES buffer for 6 h. (D) PAGE analysis: deacylation of 20 ng/µL N_21_ oligonucleotide by 10 mM Na_2_S_2_O_4_ in 50 mM HEPES buffer at 37 °C for 30 min. (E) Quantitative analysis: acylation level of N_21_ oligo when treated with three acylating reagents at 50 °C for 6 h. (F) Quantitative analysis: deacylation of N_21_ oligo by 10 mM Na_2_S_2_O_4_ in 50 mM HEPES at 37 °C for 30 min. (G) MALDI TOF analysis of **4N2CIm**-acylated and deacylated oligos. Reversible acylation: 10 µM N_21_ oligonucleotide acylated with 240 mM **4N2CIm** in 100 mM MES buffer for 6 h, then deacylated by 10 mM Na_2_S_2_O_4_ in 50 mM HEPES buffer at 37 °C for 1 h. Temp. – temperature. Avg. No. of Ac. – Average number of acylation per strand. Gels were stained with SYBR-Gold. Band intensities were calculated by Image Lab software. Data are shown as mean ± SD from at least three replicates. T-test: ns, P > 0.05; *, P ≤ 0.05; **, P ≤ 0.01; ***, P ≤ 0.001; ****, P ≤ 0.0001.

We then investigate the deacylation triggered by nitro reduction. The N_21_ oligos (20 ng/µL) were first acylated by 240 mM **4N2CIm** or **4NIm** in 100 mM MES buffer at 50 °C for 6 h. After purification via EtOH precipitation, the acylated RNAs were treated with 10 mM freshly prepared Na_2_S_2_O_4_ in 50 mM HEPES buffer (pH 7.5) at 37 °C for 30 min. The diminished upper shifted bands in lane 3 of Figure 1D (also see Figure S2) indicated the successful deacylation of **4N2CIm**-treated oligo by nitro reduction. The quantitative analysis (Figure 1F) revealed a notable decline in both the acylation percentage (from 75.8% to 37.9%) and the average number of acylated nucleotides (from 1.4 to 0.6). In contrast, deacylation of the **4NIm**-treated oligo proved inefficient within 30 minutes, showing no significant changes in either the acylation percentage or the average number of acylations. However, extending the incubation period to one hour enabled the acylation removal of the **4NIm**-treated oligo (Figure S3). We also collected MALDI-TOF mass spectra of 4**N2CIm**-acylated and Na_2_S_2_O_4_-deacylated RNA (Figure S8). The vanishing of the acylation adduct further proved the reversibility of acylation. The higher reductive sensitivity of **4N2CIm** may be explained by the lower electron density on the aromatic ring caused by the Cl atom, making the nitro group more susceptible to electron addition.^24^ Given its superior reactivity in both acylation and deacylation steps, **4N2CIm** was selected for subsequent application.

After establishing the reversibility strategy with a short oligonucleotide, we then moved to the acylation of functional RNA. We selected M1 RNA, the catalytic core of bacterial RNase P, and a model substrate, pATSerUG, to modulate the RNA-RNA interaction through reversible acylation.^25^ The substrate was labeled with Cy5 dye at its 5’-end to enable visualization of the 5’-leader degradation product. For effective substrate degradation, M1 RNA required recognition of the acceptor stem domain in the pATSerUG substrate.^26^ When the interaction between the ribozyme and the substrate was disturbed by the acylation modification due to improper folding or steric hindrance, the pATSerUG would stay intact (Figure 2A). We incubated the 25 ng/µL pATSerUG with increasing concentration of **4N2CIm**. Then, the acylated pATSerUG was incubated with the M1 RNA. The degradation product was analyzed by 12% dPAGE, and the yield was determined by the band intensities of fulllength substrates and 5’-leader. As the gel image showed (Figure 2B), the majority of the DMSO-treated (mock acylated) substrate was degraded by the ribozyme after one hour; however, the degradation yield dropped significantly when the substrate was treated with 240 mM **4N2CIm**. Similar results were observed from the degradation of the substrate when ribozyme (48 ng/µL) was acylated (Figure 2C); no 5’-leaders were detected in the lane 3-5 and 7-9. The study for the recovery of the RNA-RNA acylation by Na_2_S_2_O_4_ was followed. After the treatment of 10 mM Na_2_S_2_O_4_ for an hour, the substrate was subjected to M1 RNA degradation again (Figure 2D and S5). The degradation yield increased from 7.8 % to 82.6% after one hour of incubation with ribozyme (Figure 2E). The cleavage activity of ribozyme was also partially restored via nitro reduction, indicated by the rise of degradation yield from 48.3% to 68.4% after Na_2_S_2_O_4_ treatment (Figure 2F, 2G, and S6). By the RNA degradation assay, our acylating reagent was proven to be an effective tool for RNA-RNA interaction modulation and interrogation.

**Figure 2.**
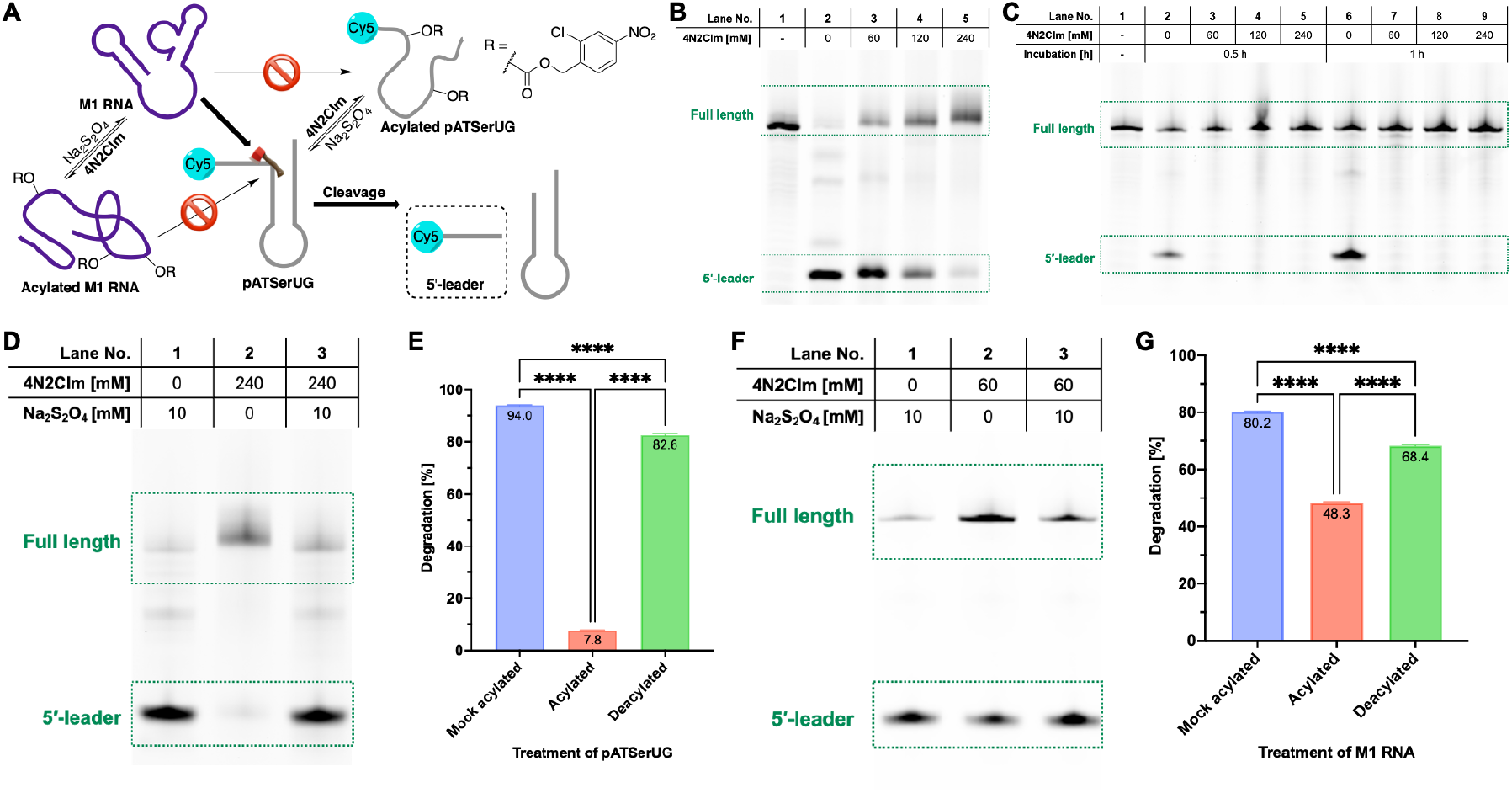
Control of RNA-RNA interaction by reversible acylation. (A) Scheme of pATSerUG degradation by M1 RNA. The acylated pATSerUG should not be degraded, and the acylated RNA should not have catalytic activity. (B) PAGE analysis: degradation of acylated pATSerUG by M1 RNA at 37 °C for 1 h. (C) PAGE analysis: degradation of pATSerUG by acylated M1 RNA at 37 °C. The 5′-leader bands were only detected in lane 2 and 6; therefore, no quantitative analysis could be done. (D) PAGE image and (E) quantitative analysis: reversible acylation of pATSerUG substrate and its degradation. (F) PAGE image and (G) quantitative analysis: reversible acylation of M1 RNA and its cleavage activity. Lane 1 (A and B) was only loaded with pATSerUG without the treatment of M1 RNA. The percentage of degradation was calculated by the Image Lab software. Data are shown as mean ± SD from three replicates. T-test: ns, P > 0.05; *, P ≤ 0.05; **, P ≤ 0.01; ***, P ≤ 0.001; ****, P ≤ 0.0001. Degradation condition: 30 ng/µL M1 RNA and 8.5 ng/µL pATSerUG were incubated in the reaction buffer (40 mM MgCl_2_, 50 mM Tris-HCl pH 7.2, 5% PEG8000, 100 mM NH_4_Cl).

In addition to RNA-RNA interactions, RNA-protein interactions (RPIs) may be regulated by reversible acylation. The steric hindrance from acylation groups impairs protein binding to RNA.^9^ To study this regulation, we employed enhanced green fluorescent protein (EGFP) mRNA as a reporter gene. When transfected into cells, acylation of the open reading frame (ORF) blocks ribosomal translation, suppressing green fluorescence (Figure 3A).^11, 23^ To test this, we treated 15 ng/µL EGFP mRNA with DMSO (mock acylation) or **4N2CIm** at 50 °C for 3 hours. After EtOH precipitation, the mRNA was split into two groups: one was incubated with 20 mM Na_2_S_2_O_4_ at 37 °C for 15 minutes to yield deacylated mRNA, while the other was incubated in HEPES buffer (mock deacylation) to maintain acylation. Both mRNA samples were then transfected into HeLa cells using Lipofectamine. The intensities of green fluorescence from these cells matched our expectations: cells transfected with DMSO-treated mRNA showed the strongest, those with acylated mRNA the weakest, and those with deacylated mRNA in between (Figure 3B). Quantitative analysis revealed that the translation efficiency of EGFP mRNA dropped to 11.9% when acylated by 40 mM **4N2CIm** and 0.9% when acylated by 80 mM **4N2CIm** (Figure 3C), suggesting the blockage of translation by acylation. Cells transfected with EGFP mRNA deacylated by sodium dithionite exhibit a stronger fluorescence signal, implying that the activity of the mRNA was partially recovered. We detected 53.1% (acylated by 40 mM **4N2CIm**) and 43.7% (acylated by 80 mM **4N2CIm**) relative fluorescent intensity from these cells. These results confirmed that acylation on mRNA was reversible and viable for cellular applications.

**Figure 3.**
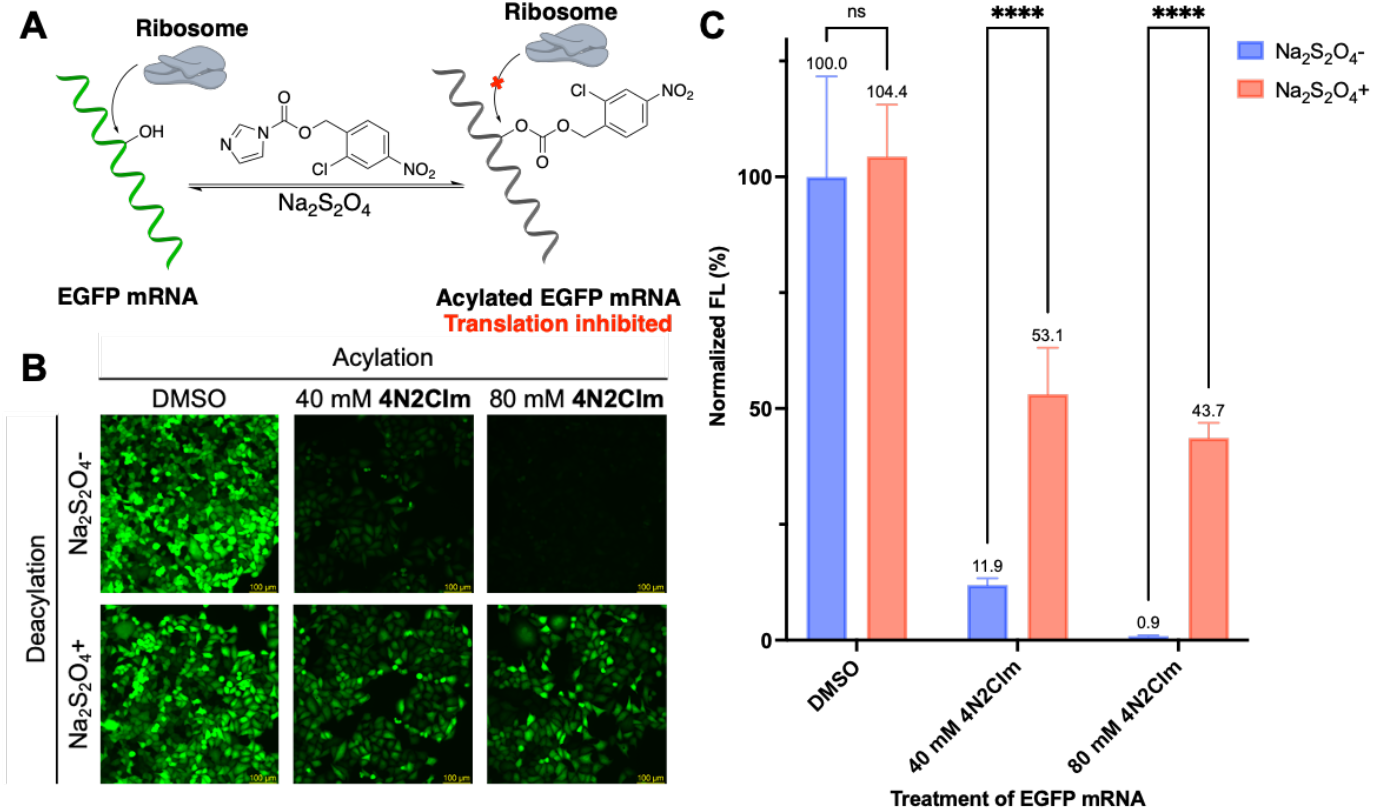
Functional control of EGFP mRNA *in vitro*. (A) Scheme of translational control of EGFP mRNA. (B) images of HeLa cells transfected with EGFP mRNA subjected to different treatments. (C) Quantitative analysis of fluorescence intensities of HeLa cells. Data are shown as mean ± SD (n=6). Scale bar: 100 µm. T-test: ns, P > 0.05; *, P ≤ 0.05; **, P ≤ 0.01; ***, P ≤ 0.001; ****, P ≤ 0.0001.

The previous assays for recovering RNA function rely on nitro reduction with Na_2_S_2_O_4_, which does not occur naturally in cells. Instead, *in cellulo* nitro reduction can be mediated by NQO1, which is overexpressed under hypoxic conditions.^13, 27^ As a preliminary test, we incubated 5 µM acylated RNA with 20 ng/µL NQO1 protein in 50 mM Tris-HCl buffer with 50 µM FAD and 1 mM NADPH at 37 °C overnight. PAGE analysis revealed the disappearance of shifted bands, thereby validating the nitro reduction activity of NQO1 *in vitro* (Figure 4C and S7). The quantitative analysis (Figure 4D) showed a decline in both the percentage of acylated RNA (from 55.30% to 32.35%) and the number of acylated hydroxyl groups per strand (from 0.82 to 0.43). To create a hypoxic condition, we treated half of the HepG2 cells with cobalt chloride (CoCl_2_), which generates a low-oxygen-like environment by stabilizing hypoxia-inducible factor-1α (HIF-1α) and interfering with its degradation (Figure 4B).^28-30^ We first validated hypoxia in HepG2 cells by measuring NQO1 mRNA levels. Total RNA from HepG2 cells under normoxia or hypoxia was extracted, converted to cDNA by reverse transcription (RT), and analyzed by quantitative polymerase chain reaction (qPCR). RT–qPCR showed increased NQO1 mRNA levels under hypoxia compared with normoxia, consistent with NQO1 overexpression (Figure 4E). EGFP mRNA (15 ng/µL) was chosen as the reporter and treated with DMSO (control) or **4N2CIm** *in vitro* at 50 °C for 2 h (Figure 4A). After transfection of these mRNAs, the fluorescence intensity of HepG2 cells was measured following overnight incubation, and the relative fluorescence intensity was used as an index of functional recovery (normalized to 100% for cells transfected with DMSO-treated mRNA). We observed higher relative EGFP fluorescence in hypoxic cells than in normoxic cells when the mRNA was acylated with 20 mM **4N2CIm** (Figure 4F and 4G). Quantitative analysis revealed 57.3% relative fluorescence under hypoxia compared with 45.6% under normoxia (Figure 4F), supporting a role for NQO1 in removing acylation from EGFP mRNA. This *in cellulo* deacylation study laid the groundwork for exploring its therapeutic potential.

**Figure 4.**
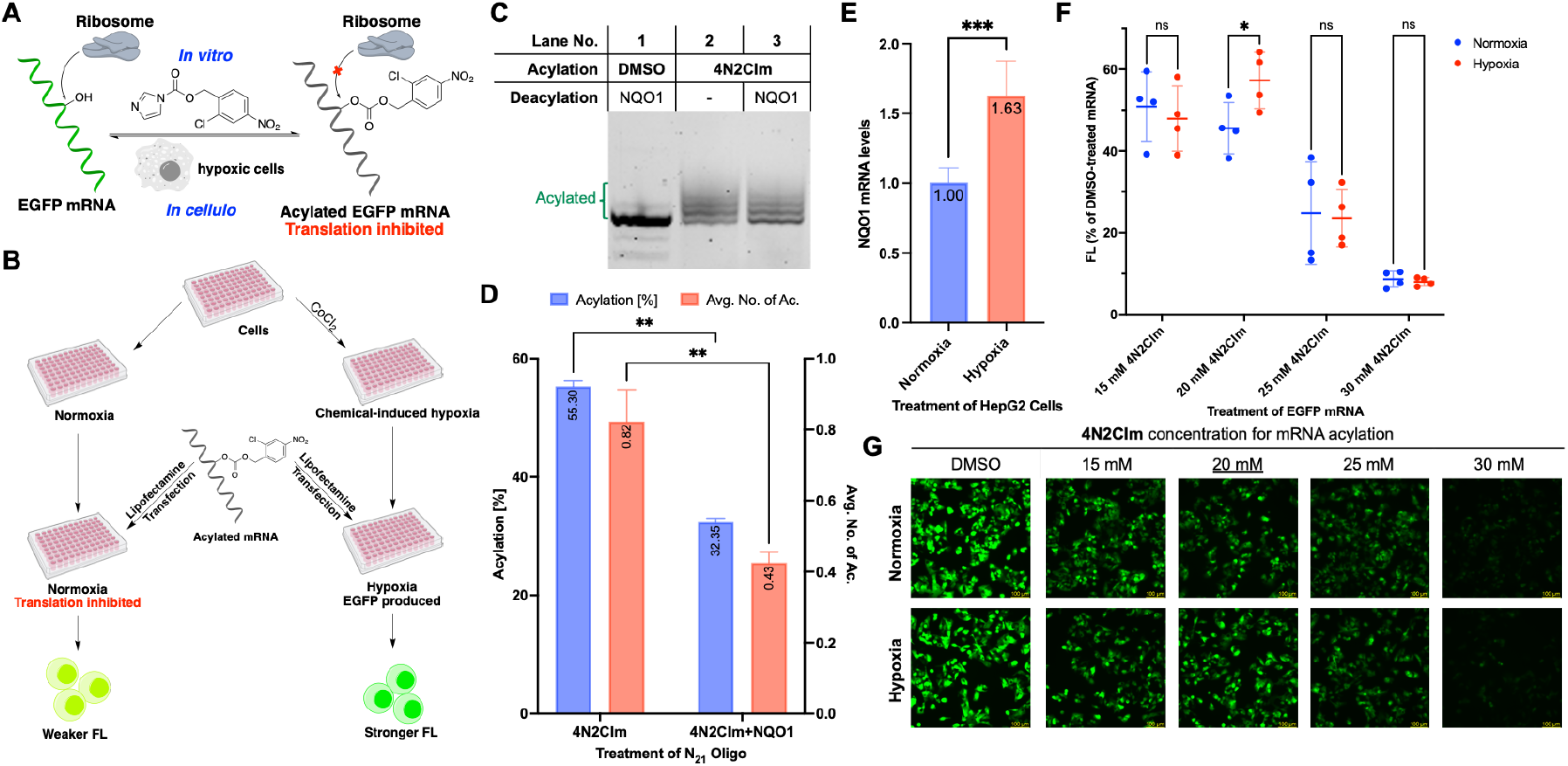
Hypoxia-induced deacylation of EGFP mRNA. (A) Scheme of *in vitro* acylation and *in cellulo* deacylation of EGFP mRNA. (B) Experimental design of hypoxia induction by CoCl_2_ and *in cellulo* deacylation in cells under hypoxia. (C) PAGE analysis: deacylation of 5 µM N_21_ oligonucleotide by 20 ng/µL NQO1 in 50 mM Tris-HCl (pH 7.4) with 100 mM NaCl and 0.5 mM EDTA at 37 °C for 18 h. (D) Quantitative analysis: acylation level of N_21_ oligo with and without NQO1 treatment. Data are shown as mean ± SD (n=4). (E) NQO1 mRNA levels were quantified by RT-qPCR using the 2^(^−^ΔΔCt) method, normalized to TBP, with normoxia set to 1. Data are shown as mean ± SD (n=5). (F) Quantitative analysis of green fluorescence of HepG2 cells. Data are presented as mean ± SD (n = 4), with individual data points shown. (G Images of HepG2 cells transfected with DMSO-treated and acylated mRNA. Scale bar: 100 µm. T-test: ns, P > 0.05; *, P ≤ 0.05; **, P ≤ 0.01; ***, P ≤ 0.001; ****, P ≤ 0.0001.

Beyond enzymatic nitro reduction, cancer cells exhibit elevated levels of other reductants, notably thiols, which are present at relatively high concentrations (millimolar) compared to other species.^31-33^ Indeed, several fluorescent probes for thiols exploit nitro group responsiveness for detection.^33^ To test the selectivity of nitro reduction, we incubated acylated N_21_ oligo with glutathione (GSH), Cysteine (Cys), and hydrogen sulfide (H_2_S). None of them induced deacylation after an hour at 37 °C (Figure S4). These findings demonstrated that nitro reduction remains unaffected by biothiols, ensuring the specificity of *in cellulo* nitro reduction processes.

In this study, we developed the **4N2CIm** acylating reagent for RNA functionality. **4N2CIm** was applied to control RNA-RNA interactions (RNAzyme degradation) and RNA-protein interactions (EGFP mRNA translation). To explore the therapeutic potential of **4N2CIm**, we chemically induced hypoxic conditions in HepG2 cells using CoCl_2_ and reactivated acylated EGFP mRNA via endogenous NQO1. This research aims to broaden the scope of chemical strategies for advancing RNA-based therapeutics, including cancer vaccines and gene therapies.

## Data availability

Details of reagents, chemical syntheses, spectral data, RNA acylation/deacylation protocols, and cell culture methods are provided in the Supplementary Information. Raw cell images are available from the corresponding author, A.M.K., upon reasonable request.

## Corresponding Author

Anna M Kietrys – Department of Chemistry, Carnegie Mellon University, Pittsburgh, Pennsylvania 15213, United States

## Author Contributions

Y.H.: data collection, experimental design, and manuscript writing/revision; K.L.: molecular design, data collection, experimental design, and manuscript writing/revision; A.K.C.: data collection and manuscript revision; C.T.: data collection; C.Y.: data collection; A.M.K.: funding acquisition, supervision, conceptualization, administration, resources, and manuscript revision.

## Acknowledgement

This research was funded by the Startup Award (MCB200159) from the Department of Chemistry at Carnegie Mellon University and supported by the Extreme Science and Engineering Discovery Environment (XSEDE) through National Science Foundation grant ACI-1548562. The funding from NSF-MRI 2117784 is highly acknowledged.

---

We developed nitro reduction-responsive acylating reagents to modulate RNAzyme-mediated pre-tRNA maturation and EGFP mRNA translation. Hypoxia-induced reactivation of mRNA in HepG2 cells highlights their potential for RNA-based therapeutics.

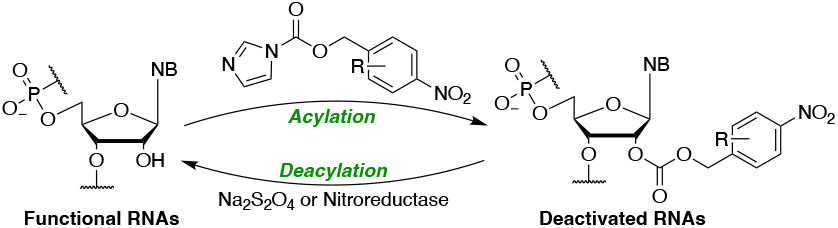
This image is for graphic abstract

