## Supporting information for "Reversible RNA Acylating Reagents with Nitro Reduction Strategy"

### 1. List of reagents and instruments

| | |
|---|---|
| **Chemical synthesis** | |
| 4-Nitrobenzyl alcohol | Combi-Blocks |
| 3-Methoxy-4-nitrobenzyl alcohol | Combi-Blocks |
| 4-Chloro-2-nitrobenzyl alcohol | Combi-Blocks |
| 1,1′-Carbonyldiimidazole | TCI |
| Methanol, HPLC grade | Fisher Scientific |
| Acetonitrile, HPLC grade | Fisher Scientific |
| Deuterated dimethyl sulfoxide (DMSO-$d_6$) | Sigma-Aldrich |
| Dimethyl sulfoxide (DMSO), anhydrous | Sigma-Aldrich |
| Ammonia | Fisher Scientific |
| Methylamine, 40% | Sigma-Aldrich |
| Triethylamine | Thermo Scientific |
| TEA • 3HF | Fisher Scientific |
| 3-Hydroxypicolinic acid | Sigma-Aldrich |
| Ammonium citrate dibasic | Sigma-Aldrich |
| Trifluoroacetic acid | Sigma-Aldrich |
| **Bioreagents** | |
| Glutathione (GSH) | Sigma-Aldrich |
| L-Cysteine (Cys) | Sigma-Aldrich |
| Sodium hydrosulfide (NaHS) | Thermo Scientific |
| Water | Corning |
| Biograde Chloroform | Sigma-Aldrich |
| Ethanol, 200 proof | Pharmco |
| Glycerol | Sigma-Aldrich |
| Orange G | Sigma-Aldrich |
| Bromophenol blue | Sigma-Aldrich |
| Xylene Cyanol FF | Sigma-Aldrich |
| Formamide | Alfa Aesar |
| MES hydrate | Sigma-Aldrich |

| | |
|---|---|
| 1.0 M HEPES buffer | Gibco |
| 1.0 M Tris-HCl buffer, pH 7.2 | Gibco |
| 1.0 M Tris-HCl buffer, pH 7.4 | Sigma-Aldrich |
| 0.5 M EDTA | Invitrogen |
| 5 M NaCl | Invitrogen |
| Azoreductase/NQO1 Protein, Human | MedChemExpress |
| Flavin adenine dinucleotide disodium salt hydrate | Invitrogen |
| NADPH tetrasodium salt hydrate | Invitrogen |
| Sodium acetate buffer solution, pH 5.2±0.1 (25 °C), for molecular biology, 3 M | Sigma-Aldrich |
| Glycogen, 5 mg/mL | Invitrogen |
| SYBR-Gold | Invitrogen |
| TEMED | National diagnostic |
| Urea | Acros Organics |
| Ammonium persulfate | Sigma-Aldrich |
| Agarose | Sigma-Aldrich |
| Acrylamide: Bis-Acrylamide 19:1 (40% Solution/Electrophoresis) | Fisher Scientific |
| Tris-Borate-EDTA, 10X Solution, Electrophoresis | Fisher Scientific |
| SequaGel – UreaGel Buffer | National diagnostic |
| SequaGel – UreaGel Diluent | National diagnostic |
| SequaGel UreaGel 19:1 Concentrate | National diagnostic |
| Cyanine 5 Phosphoramidite | Glen research |
| 2'-tBDSilyl Uridine CED phosphoramidite | Chemgene |
| 2'-tBDSilyl Adenosine (n-PAC) CED phosphoramidite | Chemgene |
| 2'-tBDSilyl Guanosine (n-PAC) CED phosphoramidite | Chemgene |

| | |
|---|---|
| 2'-tBDSilyl Cytidine (n-PAC) CED phosphoramidite | Chemgene |
| 2'-tBDSilyl Cytidine (n-PAC) 3'-lcaa CPG 500Å | Chemgene |
| $NH_4Cl$ | Alfa Aesar |
| $MgCl_2$, 1 M | Invitrogen |
| PEG8000 | Sigma-Aldrich |
| Guanosine 5'-Monophosphate Disodium Salt | Fisher Scientific |
| Phusion High-Fidelity PCR Kit | Thermo Scientific |
| DNA Clean & Concentrator-5 | ZYMO Research |
| MEGAshortscript™ T7 Transcription Kit | Invitrogen |
| CleanCap® EGFP mRNA | TriLink |
| Quick-RNA Miniprep Kit | Zymo research |
| SuperScript™ III First-Strand Synthesis System | Invitrogen™ |
| PowerTrack™ SYBR Green Master Mix for qPCR | Applied Biosystems™ |
| Dulbecco's Modified Eagle Medium (DMEM) | Gibco |
| Penicillin-Streptomycin (10,000 U/mL) | Gibco |
| Opti-MEM™ I Reduced-Serum Medium | Gibco |
| Lipofectamine MessengerMAX | Invitrogen |
| 1 × PBS buffer pH 7.4 | Gibco |
| 0.25% Trypsin-EDTA (1 ×) | Gibco |
| Fetal bovine serum, premium, USA origin | Corning |
| Sodium pyruvate | Gibco |
| Trypan blue stain | Gibco |
| Biograde DMSO | ATCC |
| **Instruments** | |
| Nuclear magnetic resonance spectrometer | Bruker Avance™ III 500 MHz and Bruker NEO 500 MHz NMR |

| | |
|---|---|
| Mass spectrometer | Thermo Scientific™ Exactive™ Plus EMR mass spectrometer |
| Confocal microscope | Leica DMi8 |
| Quantitative PCR | QuantStudio™ 5 Real-Time PCR System |
| DNA/RNA synthesizer | Mermade 4 |
| MALDI TOF | Bruker Ultraflextreme MALDI TOF/TOF MS |
| Plate reader | Tecan Spark |

### 2. Synthesis and molecular characterization

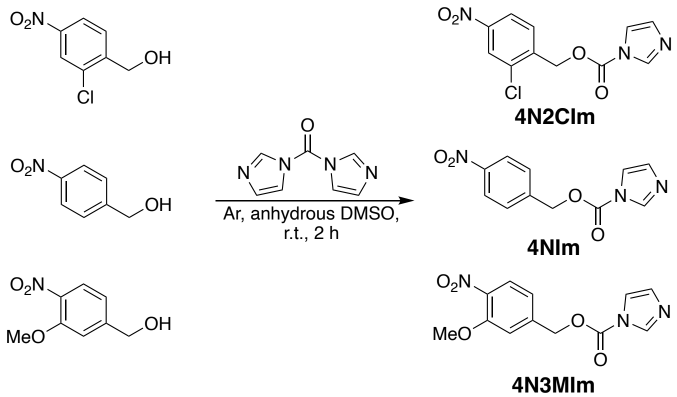


**Scheme 1.** Synthesis of acylating reagents

**General method of acylating reagent (0.4 M in DMSO) synthesis:** To a 1.5 mL Eppendorf tube, 1 eq. of nitrobenzyl alcohol (or its derivatives) was added in an Argon (Ar) environment. In a separate Eppendorf tube filled with Ar, 1.5 eq of 1,1-carbonyldiimidazole was dissolved in anhydrous DMSO (0.6 M) and quickly added to nitrobenzyl alcohol (or its derivatives) with a syringe. The reaction was mixed by vortex and incubated on a plate shaker for 2 hours at room temperature. The product in DMSO (0.4 M stock solution) was directly used as the RNA acylating reagents. For NMR data collection, the stock solution was diluted with DMSO-$d_6$.

2-chloro-4-nitrobenzyl 1H-imidazole-1-carboxylate **(4N2CIm).** $^1$H NMR (500 MHz, DMSO-$d_6$) δ 8.39 (dd, J = 3.8, 1.8 Hz, 2H), 8.25 (dd, J = 8.5, 2.3 Hz, 1H), 7.99 (d, J = 8.5 Hz, 1H), 7.70 (t, J = 1.5 Hz, 1H), 7.13 (dd, J = 1.7, 0.9 Hz, 1H), 5.63 (s, 2H). $^{13}$C NMR (126 MHz, DMSO-$d_6$) δ 148.51, 148.41, 140.00, 137.93, 133.44, 131.22, 131.01, 124.78, 122.91, 118.15, 66.19. HRMS m/z: $[M+H]^+$ Calcd 282.0287; Found 282.0275.

4-nitrobenzyl 1H-imidazole-1-carboxylate **(4NIm).** $^1$H NMR (500 MHz, DMSO-$d_6$) δ 8.36 (t, J = 1.1 Hz, 1H), 8.30 – 8.25 (m, 2H), 7.84 – 7.79 (m, 2H), 7.68 (t, J = 1.5 Hz, 1H), 7.11 (dd, J = 1.7,

0.9 Hz, 1H), 5.60 (s, 2H). $^{13}C$ NMR (126 MHz, DMSO-$d_6$) δ 148.62, 147.89, 142.84, 137.90, 130.92, 129.31, 124.13, 118.13, 68.28. HRMS m/z: $[M+H]^+$ Calcd 248.0677; Found 248.0666.

3-methoxy-4-nitrobenzyl 1H-imidazole-1-carboxylate **(4N3Mim).** $^1H$ NMR (500 MHz, DMSO-$d_6$) δ 8.36 (d, J = 1.1 Hz, 1H), 7.92 (d, J = 8.3 Hz, 1H), 7.68 (t, J = 1.5 Hz, 1H), 7.52 (d, J = 1.6 Hz, 1H), 7.29 – 7.25 (m, 1H), 7.11 (dd, J = 1.7, 0.9 Hz, 1H), 5.52 (s, 2H), 3.96 (s, 3H). $^{13}C$ NMR (126 MHz, DMSO-$d_6$) δ 152.48, 148.63, 142.09, 139.45, 137.93, 130.89, 125.71, 120.03, 118.16, 114.07, 68.44, 57.26. HRMS m/z: $[M+H]^+$ Calcd 278.0782; Found 278.0773.

### 3. Procedures for reversible acylation and *in cellulo* experiments

**DNA and RNA sequences.**

**$N_{21}$ oligonucleotide** (customized synthesis from IDT): 5′-GCA CCA UCU UCU UCA AGG AdTdT-3′

**pATSerUG RNA** (synthesized by Mermade 4 synthesizer):

5′-Cy5-GAUCUGAAUGGAGAGAGGGGGUUCAAAUCCCCCUCUCUCCGCCAC-3′

**M1 RNA:**

5′-
GAAGCUGACCAGACAGUCGCCGCUUCGUCGUCGUCCUCUUCGGGGGAGACGGGCG
GAGGGGAGGAAAGUCCGGGCUCCAUAGGGCAGGGUGCCAGGUAACGCCUGGGGG
GGAAACCCACGACCAGUGCAACAGAGAGCAAACCGCCGAUGGCCCGCGCAAGCGG
GAUCAGGUAAGGGUGAAAGGGUGCGGUAAGAGCGCACCGCGCGGCUGGUAACAG
UCCGUGGCACGGUAAACUCCACCCGGAGCAAGGCCAAAUAGGGGUUCAUAAGGUA
CGGCCCGUACUGAACCCGGGUAGGCUGCUUGAGCCAGUGAGCGAUUGCUGGCCUA
GAUGAAUGACUGUCCACGACAGAACCCGGCUUAUCGGUCAGUUUCACCU-3′

**EGFP mRNA:** CleanCap® EGFP mRNA (5moU)- (L-7201) from TriLink

**NQO1 primers:**

5′-CCTGCCATTCTGAAAGGCTGGT-3′ (forward)

5′-GTGGTGATGGAAAGCACTGCCT-3′ (reverse)

**TBP primers:**

5′-CACGAACCACGGCACTGATT-3′ (forward)

5′-TTTTCTTGCTGCCAGTCTGGAC-3′ (reverse)

**Ethanol precipitation and wash.** After the incubation, the reaction mixture was brought to 86.5 µL with water, then 9.9 µL sodium acetate (3 M, pH 5.2), 4.6 µL glycogen (5 mg/mL), and 404 µL ethanol were added. The mixture was kept at -80 °C overnight before centrifugation at 4 °C (30 min, 20600 rcf). The supernatant was removed by aspiration. Then 500 µL ice-cold 75% EtOH was added to the tube and centrifuged at 4 °C (10 min, 20600 rcf). After removing the supernatant, the pellet was air-dried for 10 min before resuspension.

**2′-OH acylation of $N_{21}$ oligo.** In a 10 µL reaction, 20 ng/µL $N_{21}$ oligo was incubated with 240 mM acylating reagents or DMSO in 100 mM MES (pH 7.5). The percentage of DMSO in the final reaction mixture was 60%. After incubating at different temperatures (37, 45, or 50 °C) for 6 hours, the reaction was mixed with 10 µL 2 × orange G in formamide (loading dye). The sample was denatured by heating the RNA at 90 °C for 3 min and immediately cooling it in an ice bath. Finally, the reaction was analyzed by 30% denaturing polyacrylamide gel electrophoresis (PAGE, monomer ratio: 19:1). The gel was visualized by the SYBR-gold channel.

**Reversible acylation of $N_{21}$ oligo by chemicals.** For acylation, 2 µL × 100 ng/µL $N_{21}$ oligo, 1 µL $H_2O$, 1 µL × 1 M MES buffer (pH 7.5), and 6 µL **4N2CIm** or **4NIm** (0.4 M in DMSO) were incubated at 50 °C for 6 hours. The reaction was then washed with biograde $CHCl_3$, and the RNA was precipitated by EtOH as described before. For deacylation, the RNA pellet was resuspended in water, HEPES buffer (pH 7.5, final concentration: 50 mM), and deacylation reagent ($Na_2S_2O_4$ or thiols, final concentration: 10 mM) to 10 µL (final RNA concentration: 20 ng/µL). After incubating at 37 °C for a specified time, the reaction was mixed with 10 µL orange G in formamide, denatured, and analyzed by 30% denaturing PAGE as previously described. Note: $Na_2S_2O_4$ solution is not stable and needs to be freshly prepared or stored aliquoted at < – 20 °C.

**Reversible acylation of $N_{21}$ oligo by NQO1.** For acylation, 1 µL × 100 µM $N_{21}$ oligo, 1 µL $H_2O$, 1 µL × 1 M MES buffer (pH 7.5), and 6 µL **4N2CIm** (0.4 M in DMSO) were incubated at 50 °C for 6 hours. The reaction was then washed with biograde $CHCl_3$, and the RNA was precipitated by EtOH as described before. For deacylation, the RNA pellet was resuspended in 8 µL water, 4 µL 5 × Tris-HCl buffer (pH 7.4, 250 mM Tris-HCl, 500 mM NaCl and 2.5 mM EDTA), 1 µL RNaseOut, 1 µL FAD (1 mM), 4 µL NQO1 (100 ng/ µL), and finally 2 µL NADPH (10 mM).

After incubating at 37 °C for 18 h, the reaction was mixed with 20 μL orange G in formamide, denatured, and analyzed by 30% denaturing PAGE as previously described. Note: NADPH had to be prepared as fresh and needs to be added last so that FAD can bind to the NQO1.

**MALDI TOF analysis of $N_{21}$ oligo.** The matrix was prepared by mixing 9 volumes of saturated 3-hydroxypicolinic acid in TA50 (water/acetonitrile = 1:1 with 0.1% TFA) with 1 volume of 100 mg/mL ammonium citrate dibasic in water. 10 μM $N_{21}$ oligonucleotide acylated with 240 mM **4N2CIm** in 100 mM MES buffer for 6 h, then deacylated by 10 mM $Na_2S_2O_4$ in 50 mM HEPES buffer at 37 °C for 1 h. The RNA pellet needed to be washed with 500 mL 75% EtOH twice and 500 mL 100% EtOH once after EtOH precipitation to remove the salt. Add 1 μL of matrix to the plate and let it dry in the air. Then, 1 μL of RNA in water was added to the matrix crystal and allowed it dry in the air. The addition of RNA was repeated until the desired amount of RNA was added to the plate. We added 100 pmol RNA in our experiment.

**Production of M1 RNA.** DNA template corresponding to the M1 RNA sequence with T7 promoter sequence at the 5′ end was purchased from TWIST Biosciences, amplified using PCR, and purified using DNA Clean & Concentrator-5. MEGAshortscript T7 Transcription Kit was used for RNA synthesis. 375 mM Guanosine Monophosphate (GMP) was added to the M1 synthetic reaction. This allowed the incorporation of monophosphate at the 5′ end. The reaction was carried out at 37°C for 15 hours followed by DNase I treatment at 37°C for 15 minutes. The RNA was incubated in 0.3 M NaOAc and 0.2 mg/mL glycogen at -80 °C overnight, followed by centrifugation at 20,600 rcf for 25 minutes. The pellet obtained was washed with 500 uL of 75% ethanol, and centrifuged at 14,000 rcf for 5 minutes. The supernatant was removed, and the pellet was air-dried. M1 RNA was then analyzed using a 6% denaturing polyacrylamide gel.

**Synthesis of pATSerUG.** Model pre-tRNA, pATSerUG, was synthesized using solid-support oligonucleotide synthesis at 1 μM scale with DMT-ON using a Mermade 4 synthesizer. The RNA was cleaved from the bead support, and bases were deprotected in a single step using 1100 μL 1:1 ammonia/methylamine by shaking at room temperature for 4 hours. This solution was filtered using Pierce columns, and the beads were washed twice with 300 μL water. Ammonia was evaporated under vacuum and the solution was lyophilized to obtain 2′ protected DMT-ON RNA. This was dissolved in 125 μL of DMSO and incubated in 60 μL of TEA and 75 μL of TEA/H3F to deprotect the 2′ hydroxyls. RNA was then purified, and DMT was cleaved off using an RNA purification cartridge. The obtained RNA solution was lyophilized and redissolved in nuclease-free water. The obtained RNA was analyzed using 12% denaturing polyacrylamide gel.

***In vitro* acylation of pATSerUG.** In a 10 µL reaction, 25 ng/µL pATSerUG was incubated with **4N2CIm** (60 mM to 240 mM) or DMSO, with the final DMSO percentage to be 60%. After incubating at 50 °C for 3 hours, the reaction was washed with biograde $CHCl_3$, and the RNA pellet was collected by EtOH precipitation. The air-dried RNA pellet was finally suspended in water for the RNA degradation assay.

***In vitro* acylation of M1 RNA.** In a 10 µL reaction, 48 ng/µL M1 RNA was incubated with **4N2CIm** (60 mM to 240 mM) or DMSO, with the final DMSO percentage to be 60%. After incubating at 50 °C for 3 hours, the reaction was washed with biograde $CHCl_3$, and the RNA pellet was collected by EtOH precipitation. The air-dried RNA pellet was finally suspended in water for the RNA degradation assay.

***In vitro* reversible acylation of pATSerUG.** For acylation, in a 10 µL reaction, 25 ng/µL pATSerUG was incubated with 240 mM **4N2CIm** or DMSO, with the final DMSO percentage to be 60%. After incubating at 50 °C for 3 hours, the reaction was washed with biograde $CHCl_3$, and the RNA pellet was collected by EtOH precipitation. The air-dried RNA pellet was then suspended in 44 µL water, 5 µL HEPES buffer (0.5 M, pH 7.5), and 1 µL 500 mM $Na_2S_2O_4$. After incubating at 37 °C for 1 hour, the reaction was quenched by EtOH precipitation. The air-dried RNA pellet was finally suspended in water for the RNA degradation assay.

***In vitro* reversible acylation of M1 RNA.** For acylation, in a 10 µL reaction, 48 ng/µL M1 RNA was incubated with 60 mM **4N2CIm** or DMSO, with the final DMSO percentage to be 60%. After incubating at 50 °C for an hour, the reaction was washed with biograde $CHCl_3$, and the RNA pellet was collected by EtOH precipitation. The air-dried RNA pellet was then suspended in 44 µL water, 5 µL HEPES buffer (0.5 M, pH 7.5), and 1 µL 500 mM $Na_2S_2O_4$. After incubating at 37 °C for 1 hour, the reaction was quenched by EtOH precipitation. The air-dried RNA pellet was finally suspended in water for the RNA degradation assay.

**Cleavage of pATSerUG substrate by M1 RNA.** In a 2 μL reaction, 30 ng/μL M1 RNA and 8.5 ng/μL pATSerUG were incubated in the reaction buffer (40 mM $MgCl_2$, 50 mM Tris-HCl pH 7.2, 5% PEG8000, 100 mM $NH_4Cl$). After incubating at 37 °C for 30 min or an hour, the reaction solution was mixed with 18 μL orange G in formamide, denatured, and analyzed by 12% denaturing PAGE (monomer ratio: 19:1). The gel was visualized by the Cy5 channel.

***In vitro* reversible acylation of EGFP mRNA.** For acylation, in a 10 μL reaction, 15 ng/μL EGFP mRNA was acylated by **4N2CIm** (40 mM or 80 mM) or DMSO in 100 mM MES buffer (pH 7.5), with the final DMSO percentage to be 60%. After incubating at 50 °C for 3 hours, the reaction was then washed with biograde $CHCl_3$, and the RNA was precipitated by EtOH as described before. For deacylation, in a 10 μL reaction, the air-dried RNA pellet was incubated with 20 mM $Na_2S_2O_4$ in 100 mM MES buffer (pH 7.5) at 37 °C. After incubating for 15 minutes, the reaction was quenched by EtOH precipitation. The air-dried RNA pellet was finally suspended in water (10 ng/μL) for transfection.

**Cell culture.** HeLa cells were cultured with DMEM culture medium (10% FBS, 1% Penicillin/Streptomycin). HepG2 cells were cultured with DMEM culture medium (15% FBS, 1% Penicillin/Streptomycin). Both cells were incubated at 37°C and 5% $CO_2$ with 95% humidity. For subculture, the old medium was removed when cells reached 90% confluency. After washing with PBS, cells were dissociated by trypsin-EDTA (0.25%). Trypsinization was quenched by adding fresh culture medium, and a quarter of the cells were transferred to a new vessel.

**Transfection of EGFP mRNA for *in vitro* reversible study.** 10 k/well HeLa cells were seeded in a 96-well plate with DMEM culture medium (10% FBS, 1% Penicillin/Streptomycin) and incubated overnight at 37°C and 5% $CO_2$ with 95% humidity. Before the transfection, the old media was replaced with 100 μL fresh DMEM medium. For each well, 10 μL OptiMEM medium was mixed with 0.4 μL Lipofectamine MessengerMAX and incubated for 15 min at room temperature. 50 ng EGFP mRNA in 5 μL RNase-free water was diluted with 10 μL OptiMEM medium and incubated at room temperature for 5 min. The mRNA/OptiMEM and Lipofectamine/ OptiMEM mixtures were then combined and incubated for 10 min at room temperature. The final mixture was added dropwise into each well. After incubating overnight, the cells were imaged in the phase and GFP channels of the Leica DMi8 confocal microscope (GFP excitation wavelength: 450-490 nm, emission wavelength: 500-550 nm). For quantitative analysis, the area occupied by the cells in each image was calculated using the software ilastik (pixel classification) and CellProfiler. The fluorescence intensity in each image was calculated by CellProfiler. Cells without mRNA transfection were measured as blank. The ratio of fluorescence intensity to the cell-occupying area was plotted (blank deducted).

***In vitro* acylation of EGFP mRNA for *in cellulo* deacylation study.** In a 10 µL reaction, 15 ng/µL EGFP mRNA was acylated by **4N2CIm** or DMSO in 100 mM MES buffer (pH 7.5) with the final DMSO percentage to be 60%. After incubating at 50 °C for 3 hours, the reaction was then washed with biograde $CHCl_3$, and the RNA was precipitated by EtOH as described before. The air-dried RNA pellet (150 ng) was finally suspended in 9 µL water for transfection.

***In cellulo* deacylation of EGFP mRNA.** 15 k/well HeLa cells were seeded in a 96-well plate with DMEM culture medium (15% FBS, 1% Penicillin/Streptomycin) and incubated overnight at 37°C and 5% $CO_2$ with 95% humidity. The old medium was removed when the cells reached 60 % confluency. Half of the cells were supplemented with pyruvate-free DMEM culture medium containing 100 µM $CoCl_2$, while the other half were supplemented with regular DMEM culture medium without $CoCl_2$. The mRNA was transfected into HepG2 cells after 24 h. For each well, 25 ng DMSO-treated or **4N2CIm**-acylated mRNA was transfected into cells by Lipofectamine MessengerMAX. For each well, 3.75 µL OptiMEM medium was mixed with 0.15 µL Lipofectamine MessengerMAX and incubated at room temperature for 15 min. 25 ng EGFP mRNA in 1.5 µL RNase-free water was diluted with 5 µL OptiMEM medium and incubated at room temperature for 5 min. The mRNA/OptiMEM and Lipofectamine/OptiMEM mixtures were then combined and incubated for 10 min at room temperature. The final mixture was added dropwise into each well. 4 hours later, the medium was replaced with 100 µL regular fresh DMEM medium. After incubating overnight, the cells were imaged in the phase and GFP channels of the Leica DMi8 confocal microscope (GFP excitation wavelength: 450-490 nm, emission wavelength: 500-550 nm). For quantitative analysis, the fluorescent intensities were measured by a plate reader (ex 488 nm; em 509 nm). Cells without mRNA transfection were measured as blank.

**RT-qPCR analysis of NQO1 mRNA.** 250k/well of HepG2 cell were seeded in a 24-well plate with DMEM culture medium (15% FBS, 1% Penicillin/Streptomycin) and incubated overnight at 37°C and 5% $CO_2$ with 95% humidity. On the second day, half of the cells were treated with pyruvate-free medium with 100 µM $CoCl_2$ for 24 h. Total RNA was extracted from cells using the Quick-RNA Miniprep Kit according to the manufacturer's protocol. From each sample, 900 ng of RNA was reverse-transcribed to synthesize cDNA using the SuperScript™ III First-Strand Synthesis System in a 20 µL reaction volume. The resulting cDNA was diluted 100-fold, and 1 µL was used as a template for qPCR of 10 µL. The reactions were performed with PowerTrack™ SYBR Green Master Mix (conditions shown in the table below). The analysis quantified the expression of NQO1 relative to the endogenous housekeeping gene, TBP. All experiments were conducted with five biological replicates, and each qPCR reaction was run with four technical replicates (10 µL each).

| | | |
|---|---|---|
| 95 ˚C | | 10 min |
| 40 Repeats | 95 ˚C | 15 s |
| | 60 ˚C | 60 s |
| 60 ˚C | | 60 s |
| 95 ˚C | | 15 s |
| 4 ˚C | | Forever |

### 4. Additional gel images

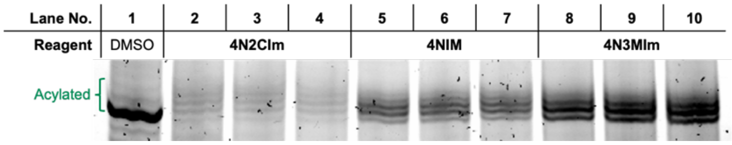


**Figure S1.** Additional repeats of $N_{21}$ acylation by three acylating reagents. 20 ng/µL $N_{21}$ oligonucleotide incubated with 240 mM acylating reagents at 50 °C in 100 mM MES buffer for 6 h.

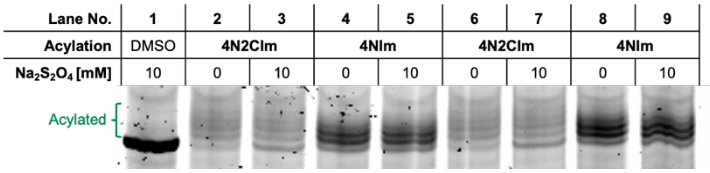


**Figure S2.** Additional repeats of $N_{21}$ deacylation by $Na_2S_2O_4$. $N_{21}$ (20 ng/µL) was acylated by 240 mM **4N2CIm** or **4NIm** at 50 °C in 100 mM MES buffer for 6 h. After EtOH precipitation, the acylated $N_{21}$ (20 ng/µL) was incubated with 10 mM $Na_2S_2O_4$ in 50 mM HEPES buffer at 37 °C for 30 min.

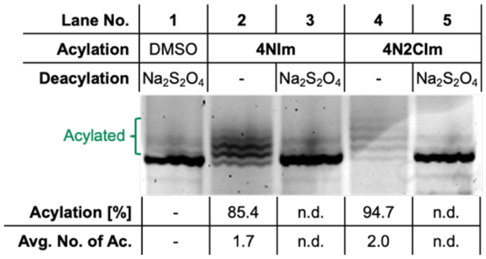


**Figure S3**. Deacylation of $N_{21}$ RNA by $Na_2S_2O_4$. $N_{21}$ (20 ng/µL) was acylated by 240 mM **4N2CIm** or **4NIm** at 50 °C in 100 mM MES buffer for 6 h. After EtOH precipitation, the acylated $N_{21}$ (20 ng/µL) was incubated with 10 mM $Na_2S_2O_4$ in 50 mM HEPES buffer at 37 °C for an hour.

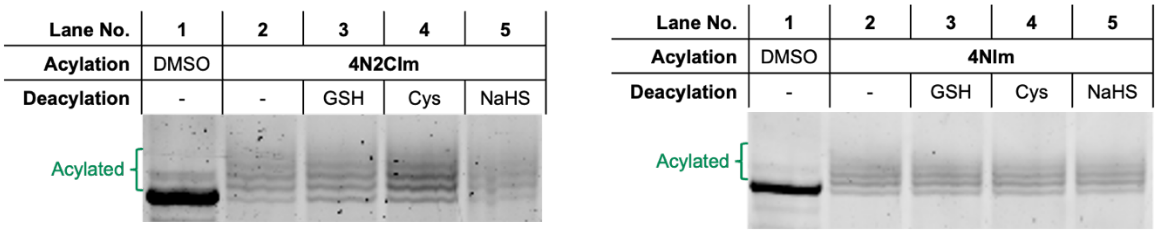


**Figure S4.** Deacylation of $N_{21}$ RNA by biothiols. $N_{21}$ (20 ng/µL) was acylated by 240 mM **4N2CIm** (left) or **4NIm** (right) at 50 °C in 100 mM MES buffer for 6 h. After EtOH precipitation, the acylated $N_{21}$ (20 ng/µL) was incubated with 10 mM biothiols in 50 mM HEPES buffer for an hour.

| Lane No. | 1 | 2 | 3 | 4 | 5 | 6 | 7 | 8 | 9 | 10 |
|---|---|---|---|---|---|---|---|---|---|---|
| 4N2CIm [mM] | - | 0 | 240 | 240 | 0 | 240 | 240 | 0 | 240 | 240 |
| $Na_2S_2O_4$ [mM] | - | 10 | 0 | 10 | 10 | 0 | 10 | 10 | 0 | 10 |

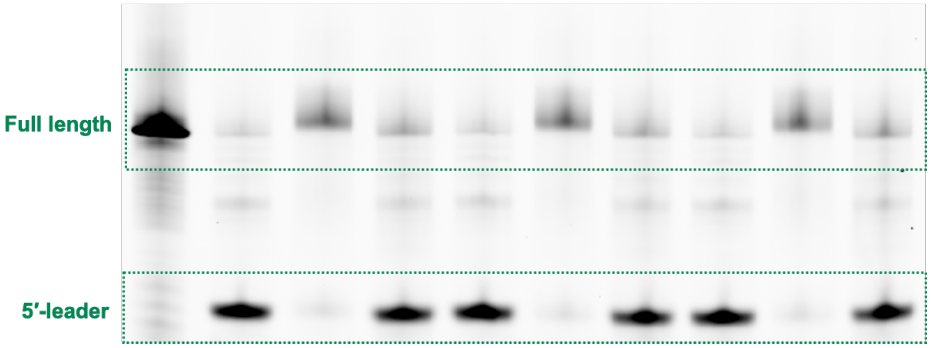


**Figure S5.** Additional repeats for the degradation of mock-acylated, acylated, and deacylated pATSerUG (8.5 ng/µL) by M1RNA (30 ng/µL) at 37 °C for an hour. Lane 1 was only loaded with pATSerUG without the treatment of M1 RNA.

| Lane No. | 1 | 2 | 3 | 4 | 5 | 6 | 7 | 8 | 9 | 10 |
|---|---|---|---|---|---|---|---|---|---|---|
| 4N2CIm [mM] | - | 0 | 60 | 60 | 0 | 60 | 60 | 0 | 60 | 60 |
| $Na_2S_2O_4$ [mM] | - | 10 | 0 | 10 | 10 | 0 | 10 | 10 | 0 | 10 |

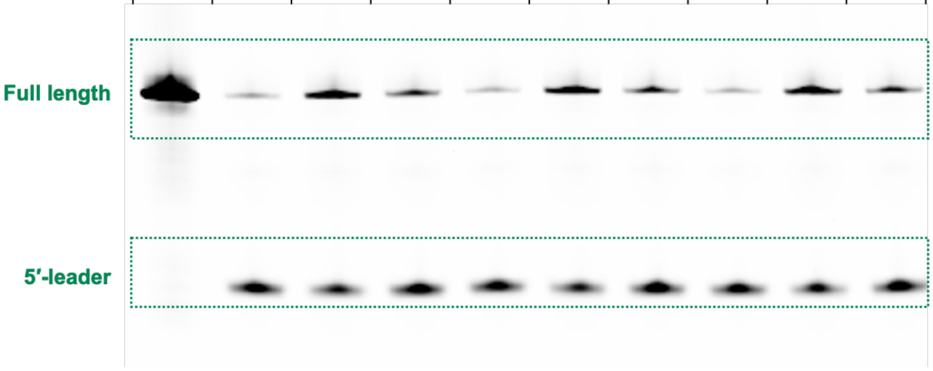


**Figure S6.** Additional repeats for the degradation of pATSerUG (8.5 ng/µL) by mock-acylated, acylated, and deacylated M1RNA (30 ng/µL) at 37 °C for an hour. Lane 1 was only loaded with pATSerUG without the treatment of M1 RNA.

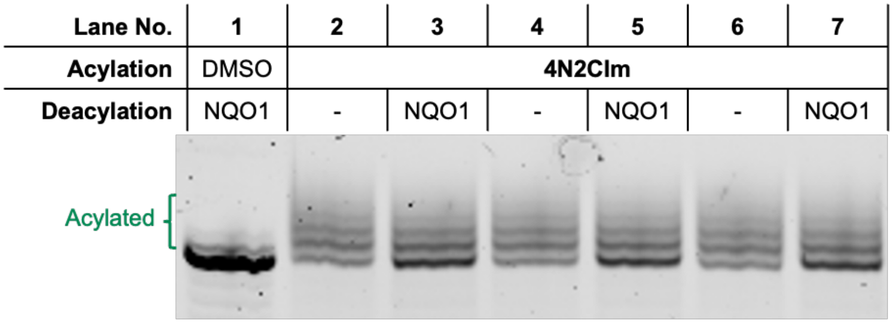


**Figure S7.** Additional repeats of $N_{21}$ deacylation by NQO1. $N_{21}$ (10 µM) was acylated by 240 mM **4N2CIm** at 50 °C in 100 mM MES buffer for 6 h. After EtOH precipitation, the acylated $N_{21}$ (5 µM) was incubated with 50 µM FAD, 1 mM NADPH and 20 ng/µL NQO1 in 50 mM Tris-HCl (pH 7.4, with 100 mM NaCl and 0.5 mM EDTA) at 37 °C for 18 h.

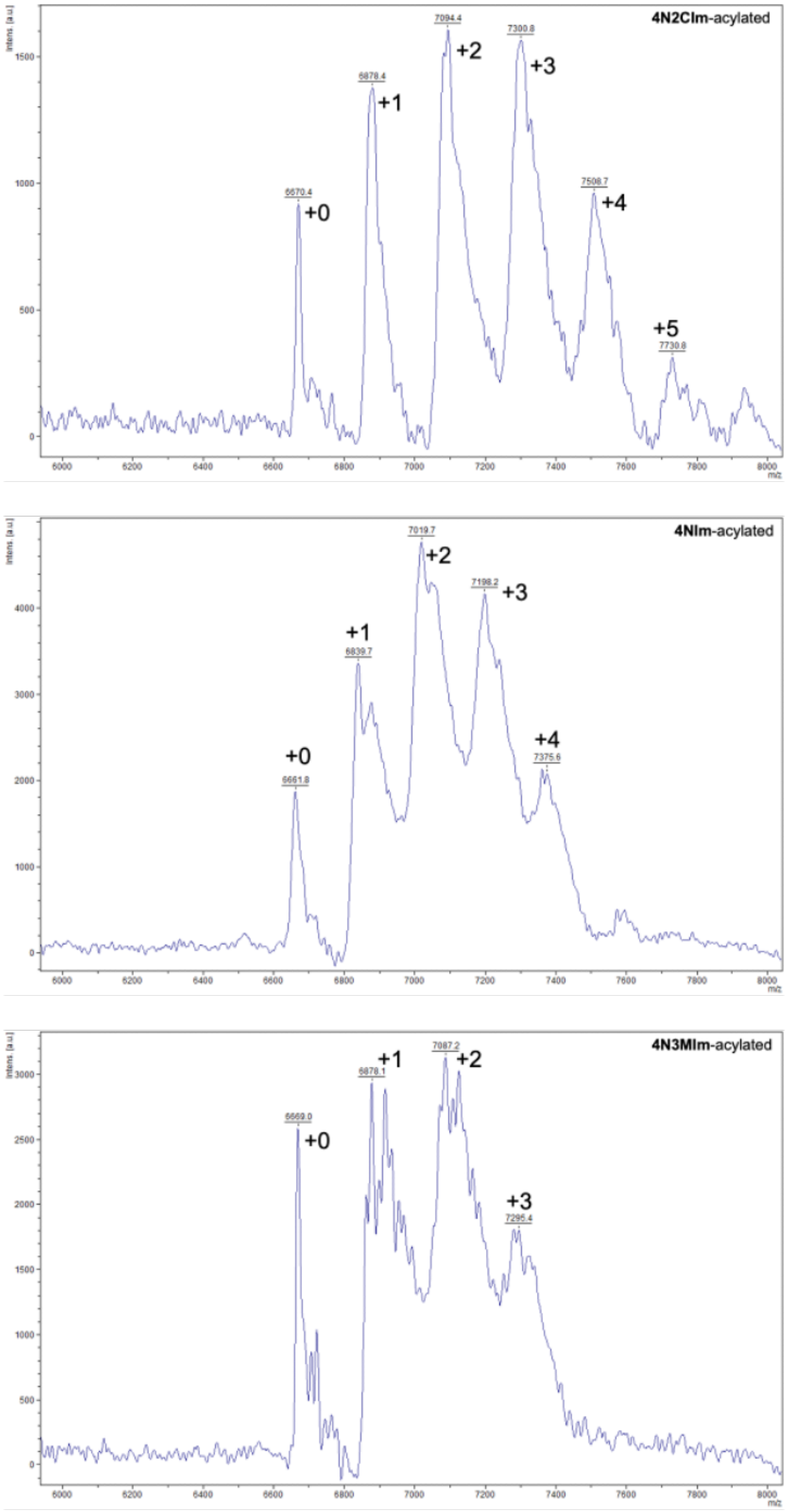


**Figure S8.** MALDI TOF analysis of acylated oligos.

### 5. Spectra

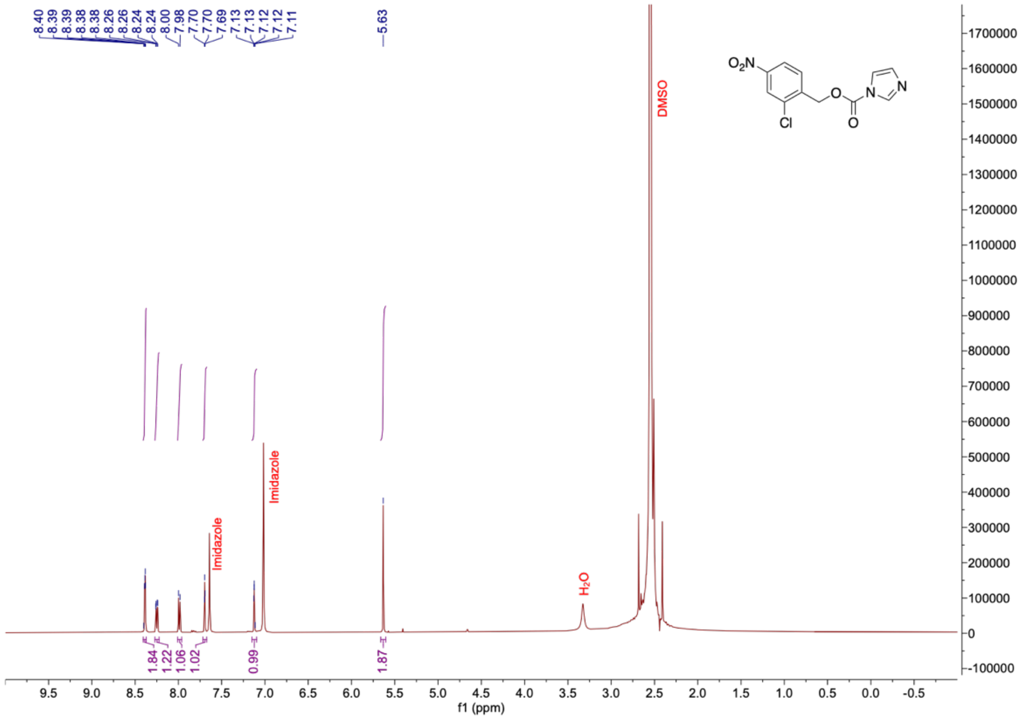


$^{1}H$ NMR spectrum of **4N2ClIm** in DMSO-$d_6$

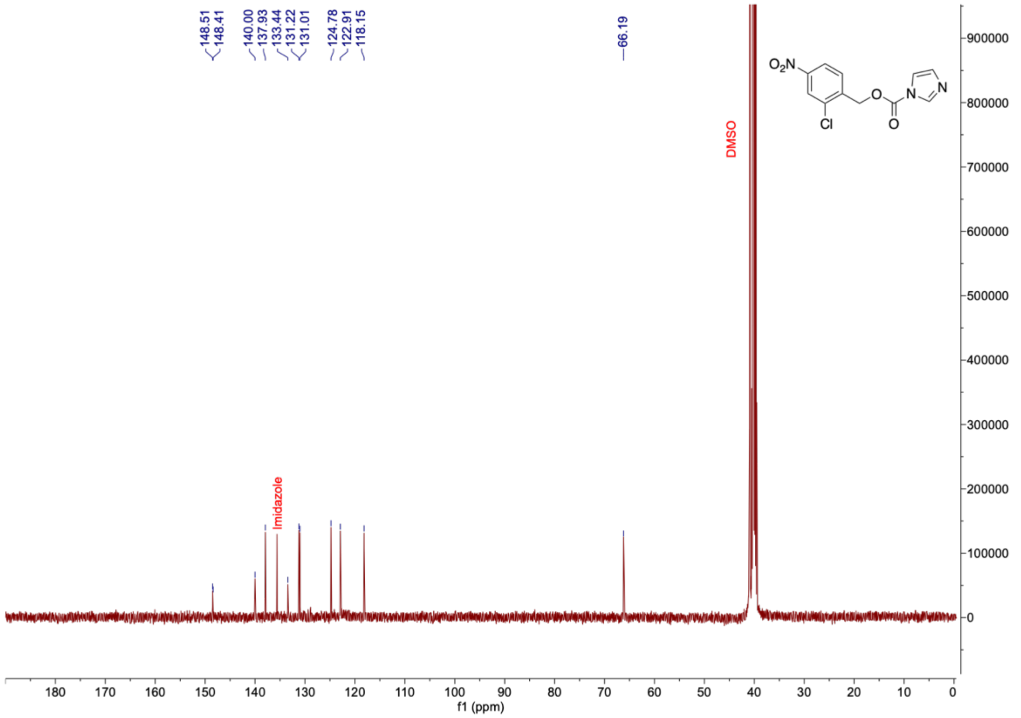


$^{13}C$ NMR spectrum of **4N2ClIm** in DMSO-$d_6$

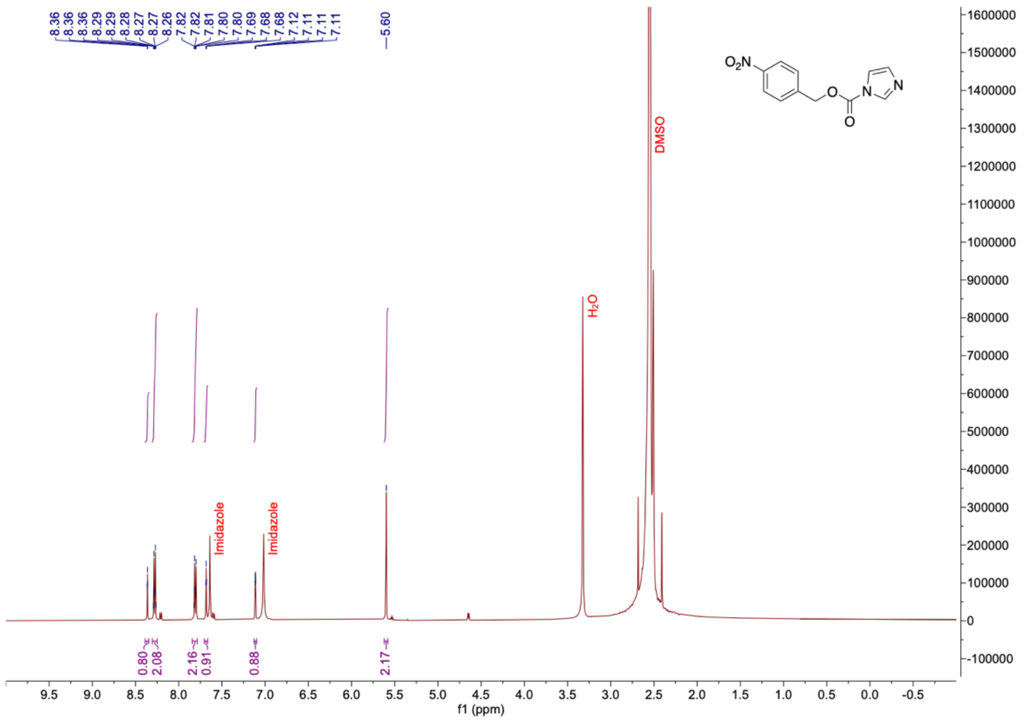


$^{1}$H NMR spectrum of **4NIm** in DMSO-$d_6$

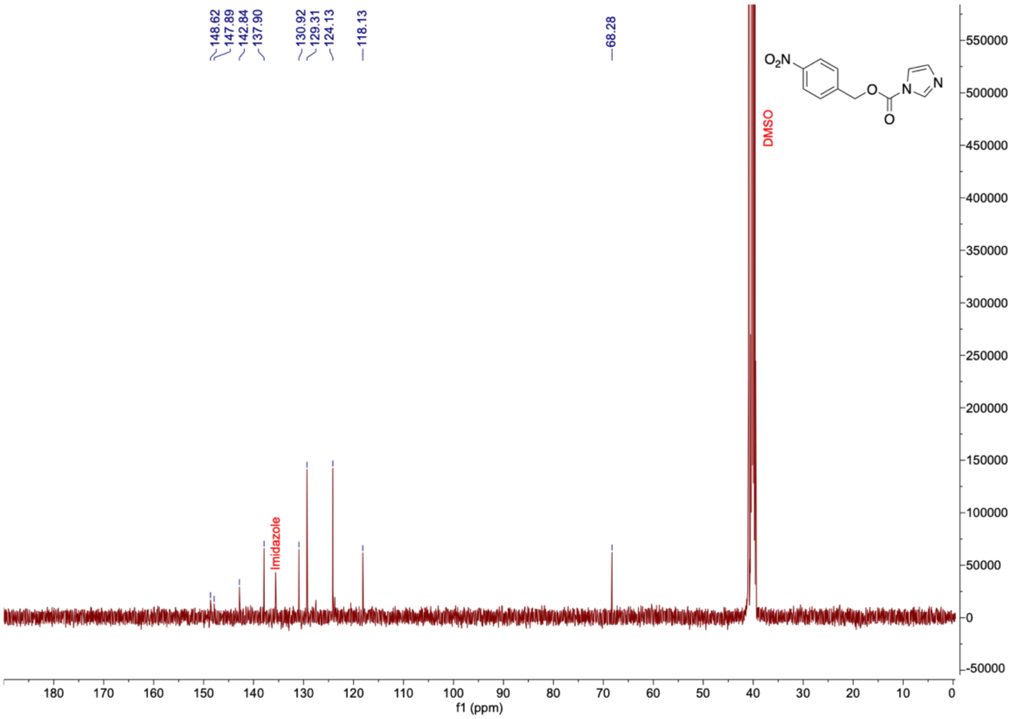


$^{13}$C NMR spectrum of **4NIm** in DMSO-$d_6$

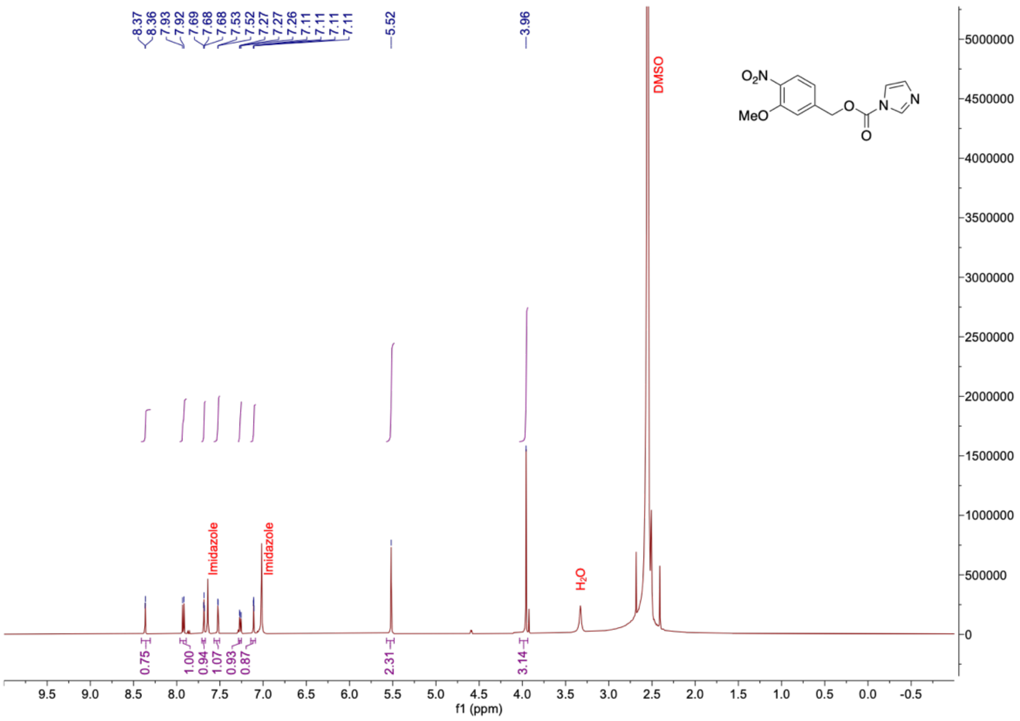


$^1H$ NMR spectrum of **4N3MIm** in DMSO-$d_6$

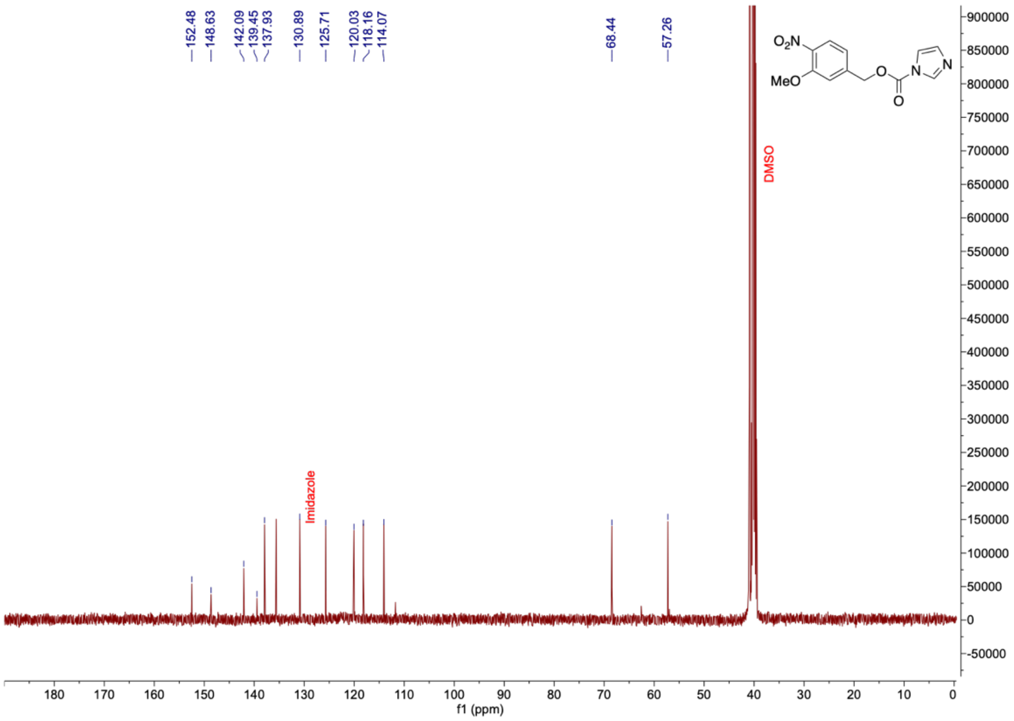


$^{13}C$ NMR spectrum of **4N3MIm** in DMSO-$d_6$

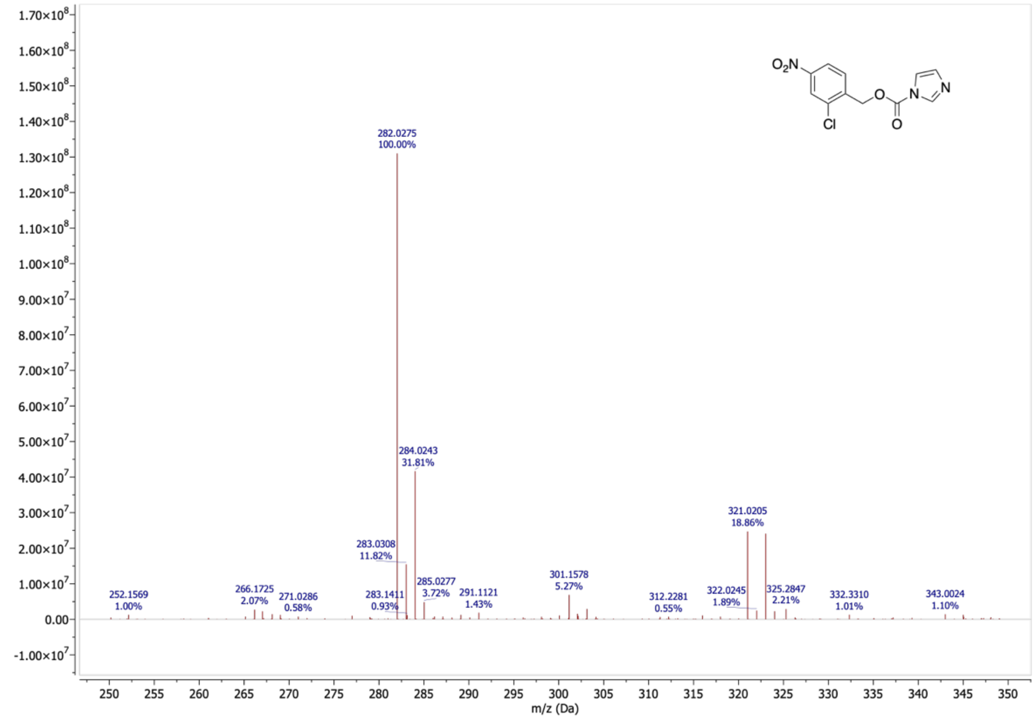


Mass spectrum of **4N2Clm**

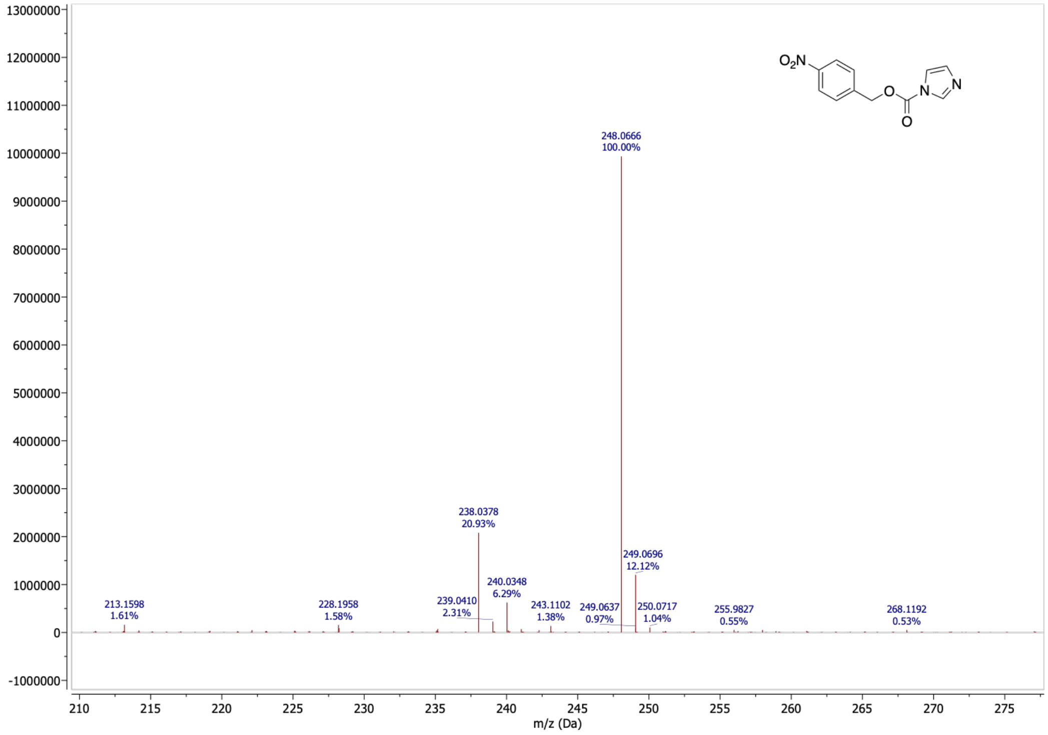


Mass spectrum of **4NIm**

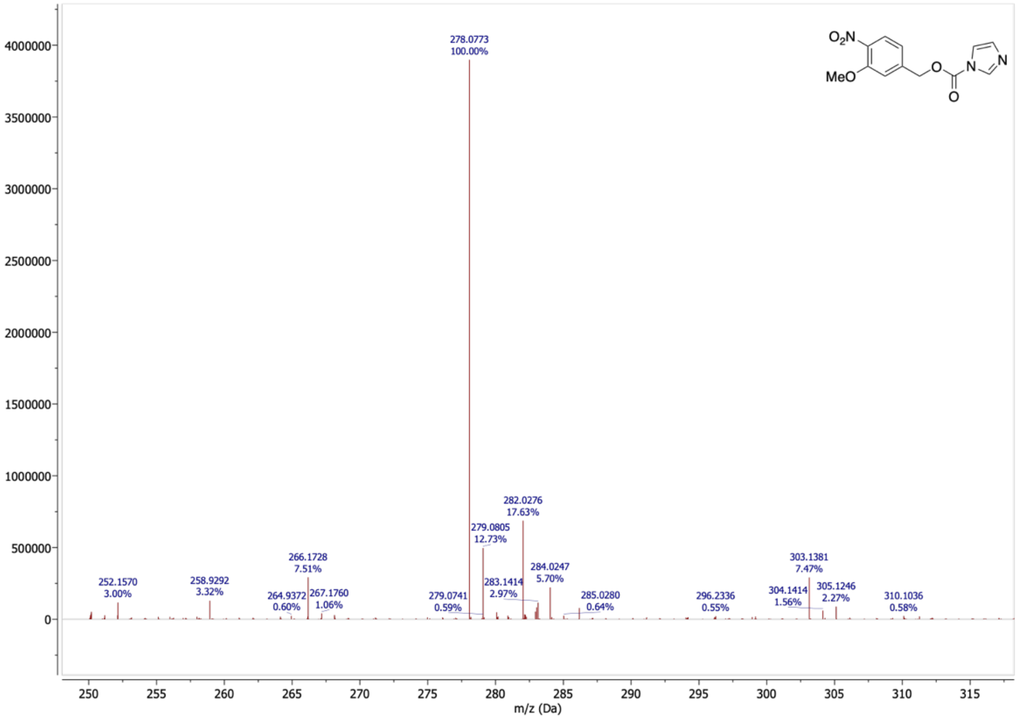


Mass spectrum of **4N3MIm**
